# Adapting to extremes: reconstructing evolution in response to changing climate over time and space in the diverse Australian plant genus *Acacia*

**DOI:** 10.1101/2021.05.08.443013

**Authors:** Xia Hua, Marcel Cardillo, Lindell Bromham

## Abstract

**Aim:** Macroevolutionary analysis is increasingly being used to study biodiversity responses to climate change, for example by using phylogenetic node ages to infer periods of diversification, or phylogenetic reconstruction of traits to infer adaptation to particular stresses. Here we apply a recently developed macroevolutionary method to investigate the responses of a diverse plant genus, *Acacia*, to increasing aridity and salinity in Australia from the Miocene to the present. We ask whether increase in tolerance of aridity and salinity coincided with periods of aridification, and if it allowed the radiation of *Acacia* into a wide range of niches.

**Taxon:** *Acacia*

**Location:** Australia

**Methods:** We applied the Niche Evolution Model (NEMo), which combines Environmental (or Ecological) Niche Modelling (ENM) with phylogenetic comparative methods (PCM) in a single statistical framework, to a large database of *Acacia* presence-only records and presence-absence survey sites in order to infer current environmental tolerances of Australia *Acacia* species and reconstruct the evolution of environmental tolerance to increasing aridity and salinity.

**Results:** We find that patterns in evolution of *Acacia*, over time and across different habitat types, are consistent with the aridification history of Australia and suggests substantial ability to adapt to high aridity and salinity.

**Main conclusions:** Our results suggest that many Acacia lineages have been able to exploit new environments created during the aridification of Australia through evolution of environmental tolerance, resulting in their current dominance of many habitats across the continent. This study demonstrates that phylogenetic studies of the evolution of responses to changing environment can move beyond application of simple trait-based models, allowing the underlying processes of speciation, adaptation and dispersal to be explicitly modelled in a macroecological and macroevolutionary context.

**Statement of significance:** *Acacia* species are found throughout Australia, from rainforests to deserts, and are striking in their environmental adaptability, so they are a perfect case study for understanding evolution of tolerance to environmental extremes in a changing climate. We use the largest database of spatial distribution records yet assembled, using both surveys and atlas data, and a new analytical method that combines the strengths of environmental niche modelling with phylogenetic comparative methods, to demonstrate rapid evolution in aridity and salinity tolerance in response to aridification of the Australian continent during the Neogene and Quaternary.

## Introduction

The climatic and environmental history of Australia during the Neogene and Quaternary Periods has been a story of increasing aridification. Dry climates may have existed from the Eocene (Carpenter et al 2014), but evidence from charcoal and pollen records reveals an increase in fire frequency and prevalence of fire-adapted, scleromorphic plant taxa, pointing to an expansion of the arid zone from the early Miocene onwards (Byrne et al 2008). As the arid zone expanded, previously widespread mesic forest habitats contracted to the coastal margins. Plant groups responded in different ways (Byrne & Murphy 2020, Crisp et al 2004, Weston & Jordan 2017). Some (such as *Nothofagus*) remained confined to humid forest conditions and contracted with these habitats to small mesic refuges. Others responded to new ecological opportunities provided by the expanding arid zone, and diversified into numerous xeromorphic lineages. Some plant families, especially Proteaceae, Myrtaceae and Fabaceae, contain both mesic and xeromorphic lineages. Even some large genera, such as *Acacia, Grevillea* and *Hakea*, have both mesic and xeromorphic representatives. Plant groups such as these, in which close relatives occupy a range of ecoclimatic conditions, provide good case studies for studying the evolution of tolerance to extreme environmental conditions.

The expansion of the arid zone in Australia led to increasingly challenging environments characterized by stressful conditions for plant growth, including drought, low soil fertility, frequent fires and high soil salinity. At the same time, areas that are extreme for most plants represent an opportunity for lineages to specialize, escape competition, and diversify (e.g. Thornhill et al 2016). Arid-adapted lineages have evolved a range of strategies to cope with water stress, broadly classified into escape, avoidance, and tolerance (Delzon 2015). All three strategies occur in *Acacia* (Moore 2013), including phyllode shedding (escape), scleromorphic structures like sunken stomata or needle-shaped phyllodes (avoidance), and physiological adaptations such as low leaf water potential (tolerance). The wide range of mechanisms adopted by *Acacia* species to cope with water stress are reflected in their distribution over extremely varied climatic conditions.

To understand how opportunities provided by drying environments shaped present-day distribution and diversity, we need to examine the dynamics of lineage adaptation in response to extreme conditions. Macroevolutionary and macroecological analyses based on phylogenies have provided a variety of ways of investigating adaptation to harsh environments (Bromham et al. 2020). For example, the ages of nodes in molecular phylogenies have been used to infer bursts of diversification associated with periods of drying in *Acacia* (Renner et al. 2020), and the movement of rainforest-adapted lineages into dry environments (Crayn et al. 2006). In addition to relative timing of diversification, phylogenies can provide information on evolution of specific adaptations to extreme conditions. For example, Crisp et al. (2011) inferred the phylogenetic history of traits such as post-fire epicormic resprouting to investigate the timing of adaptation of the eucalypts to increasing fire frequency. Jordan et al. (2008) reconstructed the evolution of stomatal protection structures (which are adaptations to dry climates) in Proteaceae to show that such structures evolved relatively few times in the ancestors of dry-climate clades. As well as physiological and morphological traits, phylogenetic approaches have been applied to reconstructing the evolution of adaptation to environmental conditions. For example, by comparing alternative evolutionary models of evolution for clades from different regions, Skeels and Cardillo (2017) showed that within four large genera (*Protea, Moraea, Banksia*, and *Hakea*), different lineages are evolving towards different climatic optima in different bioclimatic regions. Onstein et al. (2016) used the same approach to show that Proteaceae lineages in open vegetation and closed forest also have different inferred climatic optima.

Phylogenetic analyses such as these are useful, but have a number of limitations. To permit the application of standard stochastic models of trait evolution (e.g Brownian Motion (BM) or Ornstein-Uhlenbeck (OU)) in most macroevolutionary studies, the “environmental niche” is simplified to a point-estimate for a species, which is then treated as a species trait in the analysis. Usually, the trait is the mean value of an environmental variable across species’ presence locations (Evans et al 2009; Kozak & Wiens 2010; Münkemüller et al 2015; Renner et al 2020). But this approach is problematic for three main reasons.

First, the distribution of the value of environmental variables across a species’ presence locations reflects the species’ realised environmental niche (that is, the conditions under which it is found rather than the fundamental niche which describes the conditions it is able to thrive in). While the mean of the distribution may be a fair representation for narrow-range endemics found under uniform conditions, for most species an average value is not an ideal representation of the range of conditions under which the species can persist. Often the questions we want to ask are not just about the inferred midpoint of the range, but also concern the expansion into more extreme values. For example, a species with sparse presence under extreme conditions (reflected in a “long tail” of its realised niche distribution) is likely to have lower average fitness under extreme conditions than a species with frequent presence under extreme conditions (reflected in a “fat tail” of its realised niche distribution). This information can only be captured by considering the whole niche distribution, and is lost when environmental variables are summarised as average values. So we need an evolutionary model to allow changes in the whole niche distribution rather than just changes in a point-estimate of the niche distribution.

Second, even if we have an evolutionary model to allow changes in the realised environmental niche of a species over time, changes in the realised niche do not necessarily indicate evolution in the species’ fundamental niche. For a species with wide tolerance range, changes in its realised niche may be due to changes to its distribution, for example through imposition of physical barriers. We need an evolutionary model to use contemporary species distribution data to infer changes in the fundamental niche over time.

Third, standard phylogenetic methods rely on relatively simple models of trait evolution such as BM, which may not be appropriate for modelling changes in either realised or fundamental environmental niche. In particular, applying trait-evolution models to current environmental data to infer “ancestral niches” (e.g., Renner et al 2020, Skeels & Cardillo 2017) assumes that environmental traits such as aridity or temperature evolve randomly and independently along branches for each lineage. This assumption will be violated by change in response to climatic changes which apply simultaneously and directionally across all lineages within a given region. For example, Renner et al. (2020) reconstruct ancestral niches for *Acacia* by using a BM model to infer ancestral values of climatic variables such as precipitation, temperature and soil moisture. Under BM, the ancestor of a group of species found in both warm and cool environments is considered most likely to have occupied an area with a climate intermediate between the descendants, which is unlikely to be a fair model for the evolution of the Australian flora as it responds to strong directional climate change.

More fundamentally, phylogenetic analyses of evolution with changing environments compartmentalise the inference of contemporary realised niche (inferred from species distribution data) and the evolution in the fundamental niche (inferred from phylogenetic models). These are treated as two separate problems, and subject to independent analyses with separate statistical frameworks. But contemporary realised niches are the product of macroevolutionary processes: inferring these processes requires inference of the fundamental niche from contemporary distributions, so the two are fundamentally linked. The logical step, then, is to link their analyses too.

Here, we infer the history of adaptation to extreme environments by applying a recently developed method, the Niche Evolution Model (NEMo), that combines Environmental Niche Modelling (ENM) and reconstruction of the evolutionary history of fundamental niche in a single statistical framework (Hua et al. 2021). This method uses species distribution data to characterise the realised niches of each contemporary species at the tips of a phylogeny, and reconstructs evolution of the fundamental niche and the associated changes in the realised niche by modelling key driving processes: (1) shift in areas of tolerable conditions and adaptation of lineages to changing conditions (hereafter referred to as an “adaptation event”); (2) dispersal away from areas that are no longer suitable and into accessible areas of suitable habitat; (3) speciation and divergence along lineages’ environmental tolerances (hereafter “speciation event”). NEMo solves the three limitations of our current phylogenetic analyses of niche evolution listed above by modelling changes in the whole niche distribution along a phylogeny, accounting for fundamental niche and realised niche separately, and simultaneously estimating contemporary fundamental and realised niches and the evolution of fundamental niches along phylogeny.

Another strength of this method is that it allows us to infer the placement of adaptation and speciation events on the phylogeny and to reconstruct ancestral fundamental and realised niche at any time point, without making unrealistic assumptions that niches evolve under a simple stochastic model. Unlike previous approaches, NEMo can infer the occurrence density of evolutionary events on the phylogeny, allowing us to compare the timing of shifts in environmental tolerance to periods of climatic change, or compare the relative timing of evolution of tolerance along different environmental axes, using standard statistical tests, without pre-defining the covariance structure to account for phylogenetic non-independence. Information on paleoclimate can either be incorporated into the model as prior information on the occurrence rate of these events, or can be held aside as independent information against which to test the robustness of the inference (Hua et al 2021). By explicitly accounting for the effect of niche evolution history on species current niche, NEMo also improves the performance of ENMs at predicting contemporary realised niche and it can predict contemporary fundamental niche (Hua et al. 2021).

We use NEMo to investigate macroevolutionary responses of the Australian plant genus *Acacia* to aridification. *Acacia* is one of the largest and most widely-distributed plant genera in Australia, with over 900 described species occupying nearly all major habitat types, often as the dominant taxa. Many *Acacia* species are known to be tolerant of a range of extreme environmental conditions, and several previous studies have used *Acacia* species as case studies for understanding tolerance to extremes, including salinity and other aspects of soil geochemistry (Bui et al. 2014a, 2014b), and water stress (Atkin et al. 2002). There has been rapid progress in understanding the evolutionary history of *Acacia* in recent years (Maslin et al. 2003; Murphy et al. 2010; Mishler et al. 2014; Renner et al. 2020; Dale et al. 2020). Nearly all *Acacia* species are conspicuous, woody plants, so spatial presence records for *Acacia* are abundant and widespread. We combine a curated database of *Acacia* presence records with extensive presence-absence survey data, along with predictors of sampling bias, to characterise the current distribution of species in the phylogeny. We use NEMo to reconstruct the evolutionary response to increasing aridification in Australian *Acacia*, focusing on tolerance of aridity and soil salinity (Gilkes et al. 2003). Specifically, we ask two questions. First, did evolution of tolerance to increased aridity and salinity allow present-day *Acacia* species to distribute across many different habitats? Second, did increased tolerance to aridity and salinity coincide with major periods of aridification in Australia? Answering these two questions may help us understand whether *Acacia* responded to continental aridification by a burst of adaptation, allowing the genus to radiate through the arid zone.

## Methods

### 1. Phylogenetic and Spatial Data

We based our analysis on the maximum clade credibility tree of published dated phylogenies of 505 *Acacia* species, representing half of species currently recognized (Renner et al. 2020, Figure S1). To describe the current environmental conditions under which each species is found, we used a published dataset by Misher et al. (2014), which includes 132,295 carefully checked presence locations of these 505 *Acacia* species, filtered from a total of 750,000 *Acacia* presence-only records from the Australian Virtual Herbarium. To represent degree of aridity and salinity at each presence location for each species, we extracted the value of two environmental factors at each location, one from the geographic layer of maximum monthly aridity index (Williams et al. 2012) that takes into account both precipitation and evaporation and the other from the geographic layer of root zone soil electrical conductivity (Bui et al. 2017) that measures soil salinity at root zone.

Presence-only records are typically collected opportunistically, so their spatial distributions tend to be biased towards regions that are more easily accessible to researchers, especially along road networks, and areas of particular interest, such as protected areas or hotspots of diversity (Fithian et al. 2015; Haque et al. 2017), and this pattern is clearly illustrated in *Acacia* occurrence records (Figure 1). To correct for this sampling bias, we included bias factors: distance to the nearest road (using data from Coarse Cultural Topographic Datasets available at https://dev.ecat.ga.gov.au/geonetwork) and whether it is in a protected area, such as a conservation reserve or national park (from www.environment.gov.au/land/nrs/science/capad). If the parts of the species range that are accessible from roads or included in protected areas are more likely to be sampled, then we need to account for sampling bias in our model, because otherwise our sample locations represent a biased characterization of the possible conditions under which the species is found. If distance from roads or protected area are uninformative for sample location – that is, if a species range is evenly sampled and does not show more records near a road – then this parameter will have negligible influence on our results.

**Figure 1.**
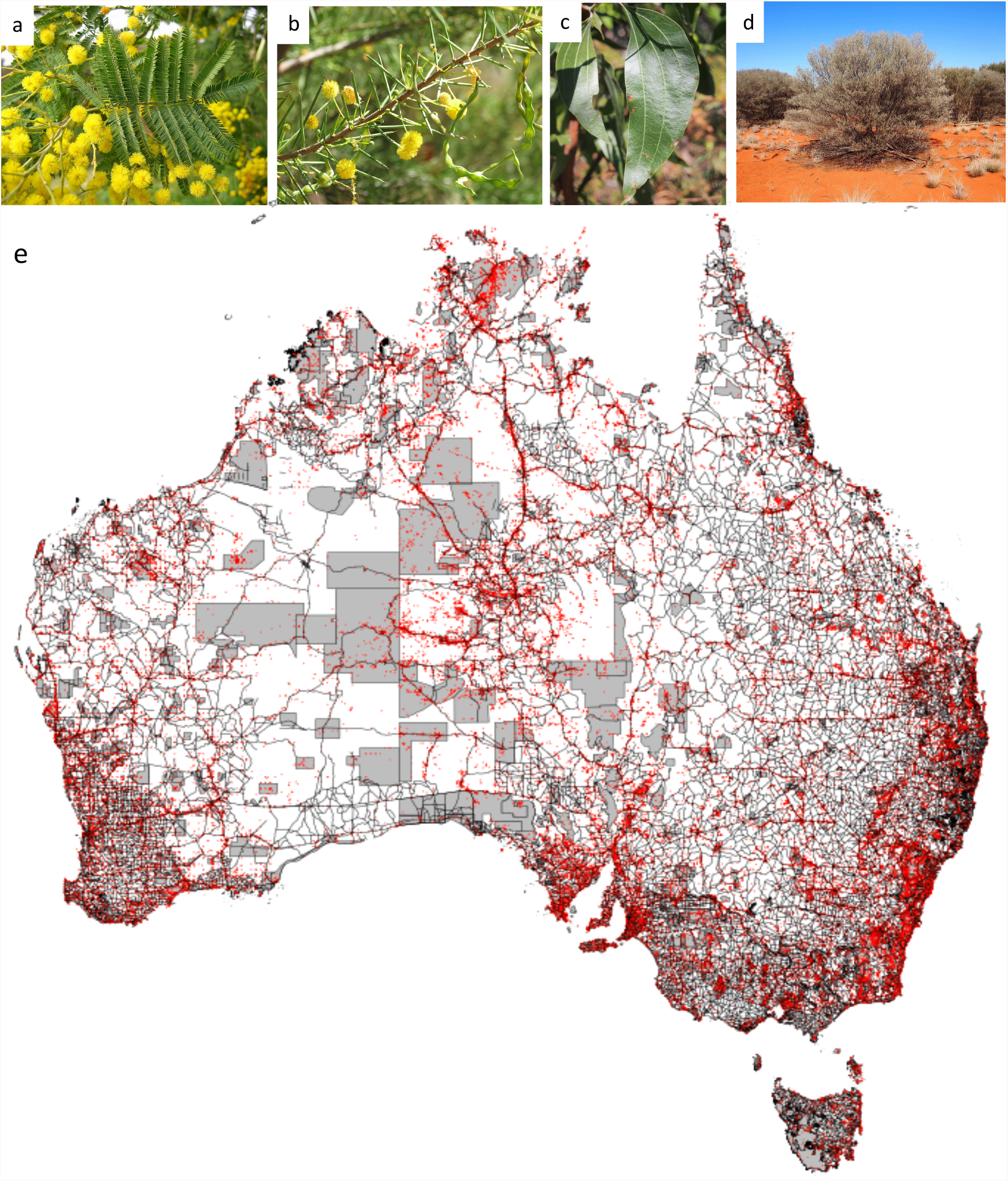
*Acacia* is a diverse genus of trees and shrubs occupying many habitats across Australia. For example, silver wattle (a, *A. dealbata*) is found in mesic habitats in southern Australia, dead finish (b, *A. tetragonophylla*) grows in arid and semi-arid areas, and yellow wattle (c, *A. flavescens*) grows in humid tropical areas. While often occurring as understory shrubs and trees, in some places *Acacia* is the dominant species, such as mulga (d, *A. aneura*). (e) Map of occurrence records illustrating the patterns of sampling bias: each red dot is a species presence location; black lines are roads in Australia; grey polygons are protected areas. Presence locations are clearly distributed along roads. For areas that are distant to roads, presence locations tend to fall in protected areas. Credit for images of *Acacia*, via Wikimedia commons under Creative Commons licenses: (a) SABENCIA Bertu Ordiales; (b) Melburnian; (c) and (d) Mark Marathon.

To improve model identifiability between the effect of environmental factors and the effect of bias factors on the presence probability of a species at a location, we also assembled species presence-absence data from government-managed floristic survey databases. Survey data is a valuable addition to herbarium records because, unlike presence-only data, it provides information on both presence and absence (i.e. places where the flora was surveyed and the species was not found). Although the location of survey sites are biased, just like the presence-only locations, the same bias applies to both the number of presences and absences at each survey site, so the number of presences is unbiased compared to the number of absences (Fithian et al. 2015). Therefore, survey data is very useful for characterising the effect of sampling bias on estimating species environmental niche (Fithian et al. 2015).

There are three government-managed online databases that allow easy extraction of the survey data and include information of each survey, including the survey sites, survey time, survey type, plot size, GPS accuracy, and custodianship. These databases are from New South Wales (BioNet Systematic Flora Survey; environment.nsw.gov.au/research/VISplot.htm), Victoria (Victorian Biodiversity Atlas flora records; environment.vic.gov.au/biodiversity/victorian-biodiversity-atlas), and Western Australia (Flora Surveys of the Yilgarn; naturemap.dbca.wa.gov.au). While these surveys include only three states (which cover nearly half of the land area of Australia), here they are only used to increase the accuracy in estimating the coefficients of the two sampling bias factors. With the information of each survey, we filtered survey data that are from complete floristic surveys conducted in the previous 15 years under government custodianship to ensure consistent standards. As a result, our presence-absence survey data included 27,850 survey sites across three states of Australia. These presence-absence data were analysed together with the presence-only locations.

### 2. Niche Evolution Model (NEMo)

NEMo uses a Bayesian framework that includes three major components: model of niche evolution, phylogenetic comparative method (PCM), and environmental niche model (ENM). NEMo models species distributions in “niche space”, not geographic space, by making assumptions about the dispersal process in the model of niche evolution (see below). We model niche space in this way because we rarely have reconstructed paleoclimate with high resolution over a large region, without which it is impossible to model species distribution over geographic space on geological time scale. So NEMo provides a widely applicable approach to allow us model the evolution of the whole niche distribution under more biologically informed processes, at the cost of making simplified assumptions on the dispersal process. In comparison, traditional PCMs make simplified assumptions on the process of niche evolution and do not model dispersal process.

The niche evolution model calculates how three classically recognized aspects of niche (fundamental, available, realized) change over time under three basic processes (adaptation, dispersal, speciation). “Fundamental niche” represents the range of conditions that each species can tolerate at a given point in time, expressed as the proportion of individuals of a species that can tolerate a given condition, or a value along a relevant environmental axis. We expect species to evolve along this environmental axis over evolutionary time, through the processes of adaptation to new conditions (by selecting for individuals at the extreme values of the distribution) and through speciation along environmental gradients (so that a species splits along its tolerance range leaving one daughter species that is more skewed toward the extreme conditions). “Available niche” describes the range of conditions in areas that are accessible to the species, regardless of whether the species can persist in those conditions. The potential of the species to occupy the available niche is governed by its fundamental niche and dispersal ability. We also include barriers to dispersal as nuisance parameters, which could represent physical or biotic barriers. “Realized niche” describes the locations where a species is actually found, which is shaped by the suitability of conditions and the opportunity to disperse to and occupy those areas.

The PCM part of the model infers the locations on the phylogeny of adaptation and speciation events, assuming these events are distributed along the branches of the phylogeny according to a Poisson process with constant occurrence rate over the phylogeny. The distribution of these events on the phylogeny gives a possible history of evolution in the fundamental environmental niche. Dispersal in niche space is modelled as a continuous process between any two events (adaptation or speciation), or nodes of the phylogeny. It is assumed to be a diffusion process to reach the realized niche at equilibrium given the potential of the species and its available niche that is truncated at dispersal barriers. So dispersal updates the realised niche as a response to the evolution in the fundamental niche and changes in the environmental conditions in the available niche. Given this history, we use the niche evolution model to calculate the three classic aspects of the niche of each contemporary (tip) species. The ENM component of the method then assess the fit of the inferred niches to the presence and/or absence locations of the tip species. The fit of the inferred niches to the data then feeds back to our inferences of the history of the evolutionary events underlying shifting environmental tolerances along the phylogeny, and the niches of tip species. Details of the method, and description of tests of its performance and reliability, are given in Hua et al. (2021). Code with step-by-step instructions for implementing the model can be found at https://github.com/huaxia1985/NEMo. Below we briefly describe the parameters of the model applied to this case study.

### 3. Accounting for sampling bias

We used an inhomogeneous Poisson point process (Renner et al. 2015) as the ENM component of NEMo. Because opportunistic sampling bias can influence the niche estimation (Fithian et al. 2015), we modified the likelihood function to account for sampling bias, using the method of Fithian et al. (2015): details are given in the Supplementary Information.

### 4. Model Parameters

NEMo has four universal parameters which are assumed to be constant across the phylogeny: heritability, dispersal rate, the occurrence rate of adaptation events, and the occurrence rate of speciation events. We apply separate NEMo analyses to the evolution of environmental tolerances in *Acacia* along aridity and salinity axes. For each of these axes, the values for the “available niche” at the root of the phylogeny were assumed to follow a normal distribution that is truncated at the observed maximum and minimum values of the axis (Williams et al. 2012; Bui et al. 2017); the values for the “fundamental niche” were assumed to follow 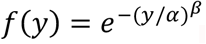, where *y* is a value along the axis (see Hua et al. 2021 for details of the function). The starting values for these parameters and their priors are described in the Supplementary Information.

### 5. MCMC approximation

For both aridity and salinity, we ran 20 independent reversible-jump Markov Chain Monte Carlo (rjMCMCs) for 3×10^6^ generations, with a thinning interval of 1000: details provided in the Supplementary Information (Figure S2). For both analyses, the 20 rjMCMCs agree on a general pattern of the amount of change in the fundamental niche along *Acacia* phylogeny (Figure S3 and S4), so we use the niche evolution history with the highest posterior probability to illustrate this general pattern for both analyses (Figure S1).

## Results

The *Acacia* phylogeny is characterized by a mixture of large, young clades (recent, rapid divergences), and smaller, older clades (longer-persisting lineages: Figure S1). The young clades, each around 10 Ma old, include most present-day species in tropical forests and grasslands and the surrounding desert areas (in clades 10 and 11; Figure S1), as well as most present-day species in subtropical forests and grasslands (in clades 7 and 8; Figure S1). The smaller, older clades, each around 20 Ma old, mainly include present-day species in habitats classified as “Mediterranean forests, woodlands & scrub” in World Wildlife Fund’s categories of terrestrial ecoregions (https://www.worldwildlife.org/biome-categories/terrestrial-ecoregions; Figure S1) (i.e., ecosystems occurring in Mediterranean type climates) and the surrounding desert areas (in clades 1, 3-6, 8; Figure S1). Present-day species in Mediterranean-type ecosystems also form smaller clades that are sisters to the recent radiations in tropical and subtropical forests and grasslands (in clades 10-11 and 7-8; Figure S1). In short, present-day species that occupy regions that are arid and prone to high salinity in today’s Australia (Mediterranean ecosystems and surrounding deserts) are scattered throughout the phylogeny, forming generally smaller clades than present-day species that occupy the relatively mesic zone with low salinity (tropical and subtropical forests and grasslands), which are found mainly in large and young clades.

### 1. The amount of niche evolution along *Acacia* lineages

The amount of evolution in the fundamental niche along the environmental tolerance axes inferred by NEMo along the phylogeny is shown as branch color in Figure S1, with red indicating increased tolerance of the extreme conditions over the branch and blue indicating decreased tolerance. Both aridity and salinity analyses infer more red than blue branches, but the result is much stronger in the salinity analysis (Figure S3 and S4), which suggests that evolutionary change, particularly on salinity axis, has been pushing species towards tolerance of more extreme conditions. We can also see this pattern in the inferred realised niche at the root of the phylogeny. The root had realised niche on aridity axis under conditions equivalent to the medium aridity level of today’s Australia, but its realised niche on salinity axis is under conditions equivalent to the lowest salinity level of today’s Australia (Figure S1). So our analyses suggest that the common ancestor of *Acacia* was likely to have been able to tolerate the aridity level of most non-desert areas in today’s Australia, but was possibly sensitive to soil salinity.

We can also compare the inferred environmental niches of present-day species in different habitats (using World Wildlife Fund’s categories of terrestrial ecoregions), as well as comparing them to the common ancestor (Figure 2). On the aridity axis, present-day species mainly found in tropical forests have the most similar environmental niches (in all three aspects: fundamental, available, realised) to the common ancestor (Figure 2A-2C), a pattern which is also reflected in the absence of niche evolution along the branches connecting the root and the most recent common ancestor (MRCA) of the radiation in tropical forests and grasslands (Figure S1). The largest changes in environmental niches (in all three aspects) are in lineages currently found in deserts, the most arid environments (Figure 2A-2C). Along the branches leading to these desert species, there are often multiple rounds of evolution to tolerate higher aridity on the phylogeny, both in the old clades and the recent radiations (Figure S1). Lineages in all the other habitats have similar inferred fundamental niche (Figure 2A), although their inferred realised niches tend to differ (Figure 2C).

**Figure 2.**
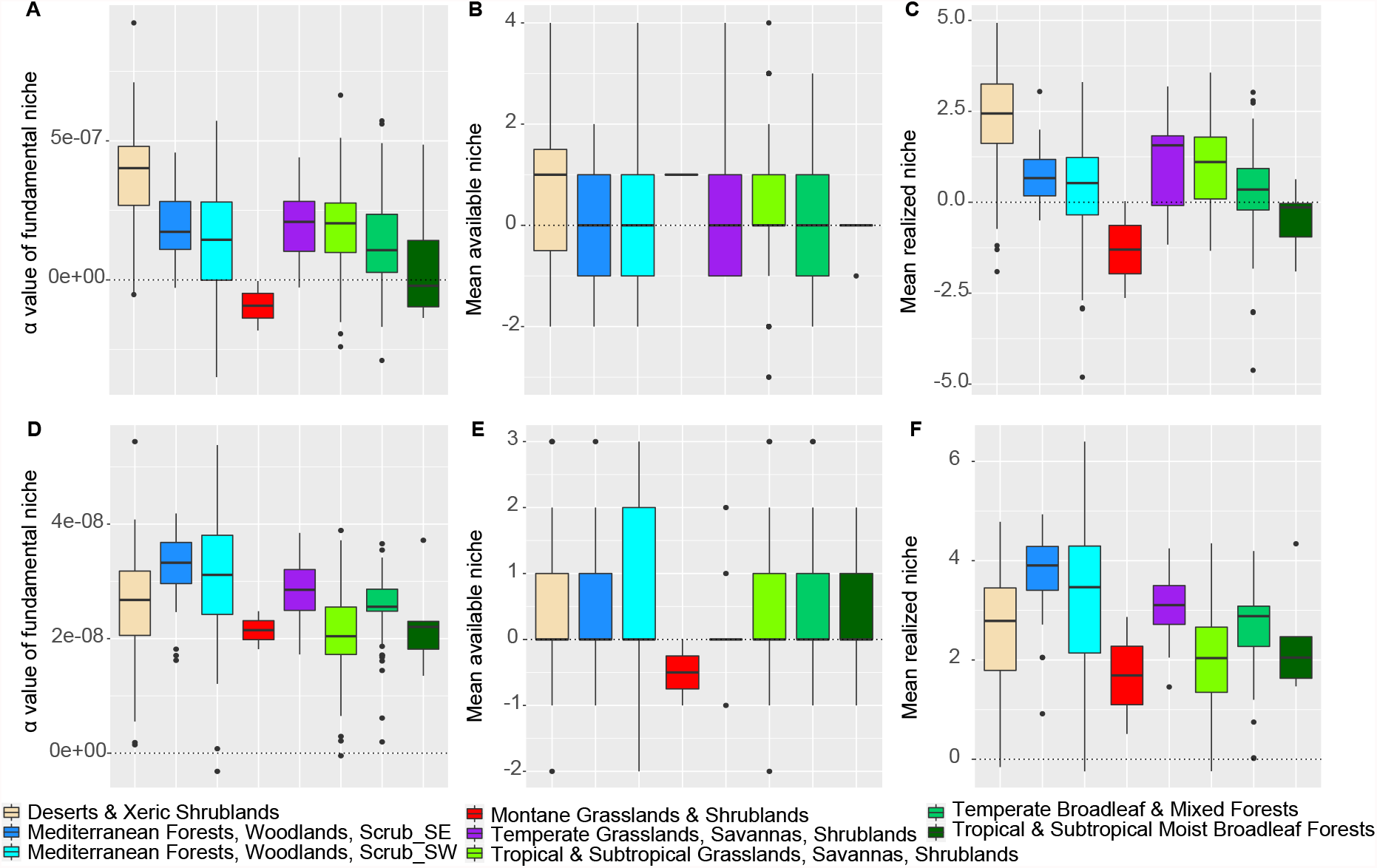
Boxplots of the total amount of change in the three aspects of environmental niche (fundamental, available, realized) from root to each tip along aridity axis (A-C) and salinity axis (D-F). Tips are grouped by the habitat that the majority of current presence location of the tip species fall in, using the same color scheme as the habitat map in Figure S1.

In contrast, on the salinity axis, our analysis inferred large shifts in both fundamental niche and realised niche towards higher salinity since the common ancestor of *Acacia* in all president-day lineages (Figure 2D-2F). The amount of shift is largest in species currently distributed in the Mediterranean ecosystems, and is smallest in extant tropical and montane species (Figure 2D-2F). Lineages in all the other habitats have similar inferred fundamental niche and realised niche (Figure 2D,2F). Shifts in most lineages are due to the major niche evolution events at the very basal branches of the phylogeny. Along the branches leading to species currently distributed in the Mediterranean ecosystems, there are multiple rounds of evolution to tolerate higher salinity, both in the old clades and the recent radiations (Figure S1).

### 2. The timing of niche evolution events

Figure S1 shows the inferred occurrences of adaptation and speciation events along the phylogeny, with red indicating events that increase the tolerance of the extreme conditions and blue indicating events that decrease the tolerance. Since there have been three significant drying periods in Australia during the Cenozoic Era: around 28-23 Million years ago (Ma), 14-5 Ma, and throughout the Pleistocene since 2.6 Ma (Owen et al. 2017), we count the number of inferred adaptation and speciation events between the time intervals of the root age, 23 Ma, 14 Ma, 5 Ma, and 2.6 Ma. A common pattern of these niche evolution events along aridity and salinity axes is that speciation events have similar occurrence density per lineage over all time intervals (Figure 3A,3C), while adaptation events have increasing occurrence density per lineage towards the present, with significantly high occurrence density during the Pleistocene (Figure 3B,3D). This burst of adaptation events along aridity and salinity axes in the Pleistocene mainly occurs in clades 7-11 that include most of the present-day species in the tropical and subtropical forests and grasslands (Figure S5).

**Figure 3.**
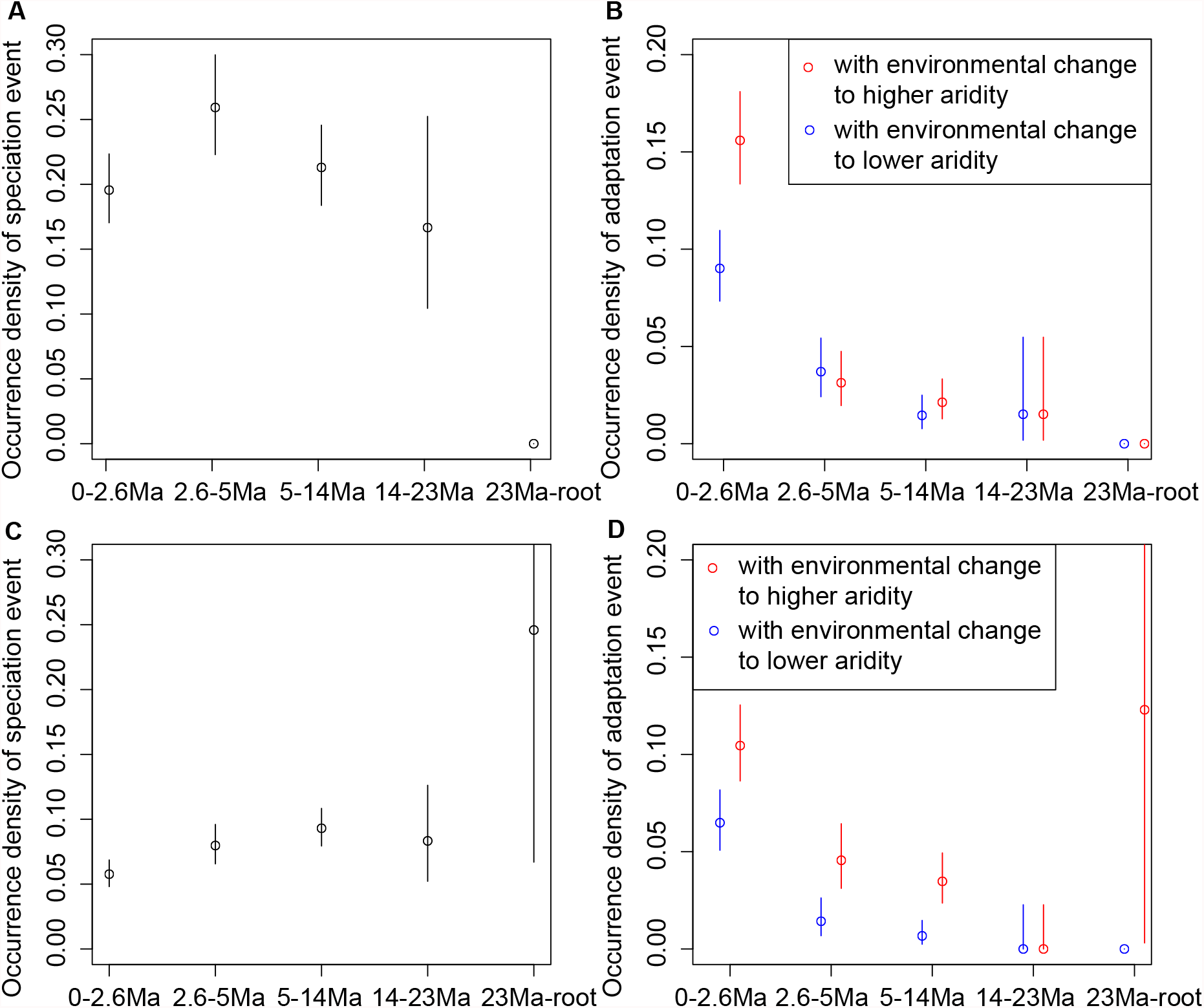
Occurrence density of the inferred speciation and adaptation events per branch that fall within each time interval. The mean of the occurrence density is calculated as the count of speciation or adaptation events occurred within a time interval divided by the sum of branch lengths within the time interval, so that the occurrence density is corrected for the number of branches. The bars show the 95% confidence interval of the occurrence density, assuming that occurrences follow Poisson process. A similar plot for each clade in the phylogeny is in Figure S5.

On the aridity axis, no adaptation or speciation events are inferred along the basal branches between the root of the phylogeny and 23 Ma (Figure S1). There tends to be higher occurrence density of adaptation events with increasing aridity than adaptation events with decreasing aridity during the drying periods in the Pleistocene and between 14-5 Ma, and vice versa between 5-2.6 Ma, the short period when Australia became wet and warm again (Figure 3B). The differences between the occurrence densities of adaptation events with increasing aridity versus decreasing aridity are statistically significant during the Pleistocene and between 14-5 Ma. Both have 95% confidence intervals of the difference in the occurrence densities above zero (the Pleistocene: 0.036∼0.096; 14-5 Ma: 0.001∼0.020). In contrast, our analysis infers continuously increased tolerance to salinity over the *Acacia* phylogeny from the very basal branches of the phylogeny to the present (Figure S1). The occurrence density of adaptation events with increasing salinity is consistently higher than the occurrence density of adaptation events with decreasing salinity over all time intervals (Figure 3D). This result is consistent with the idea that soil salt might have been continuously accumulated in Australia over geological time as a result of the antiquity and flatness of the continent, as well as the complex relationship between salinisation and aridification which is not a simple linear relationship (George et al. 2008).

### 3. Uncorrelated evolution on aridity and salinity axes

To test if evolution to tolerate aridity occurs in concert with evolution to tolerate salinity in *Acacia*, we plot the inferred amount of evolution in the fundamental niche along each branch of the *Acacia* phylogeny (i.e., the branch color in Figure S1) on the aridity axis versus that on the salinity axis in Figure 4A. The Pearson’s correlation test suggests no correlation in the amount of evolution on the aridity and salinity axes (*t*_1004_=0.47, *p*=0.64; Figure 4A). Yet, present-day species that tolerate high level of salinity tend to tolerate high level of aridity (*t*_503_=2.83, *p*=0.005; Figure 4B), which is mostly driven by species in the deserts (*t*_97_=3.72, *p <*0.001), the tropical grasslands (*t*_95_=4.23, *p <*0.001), and the Mediterranean forests in South Australia (*t*_31_=2.20, *p*=0.04; Figure S6).

**Figure 4.**
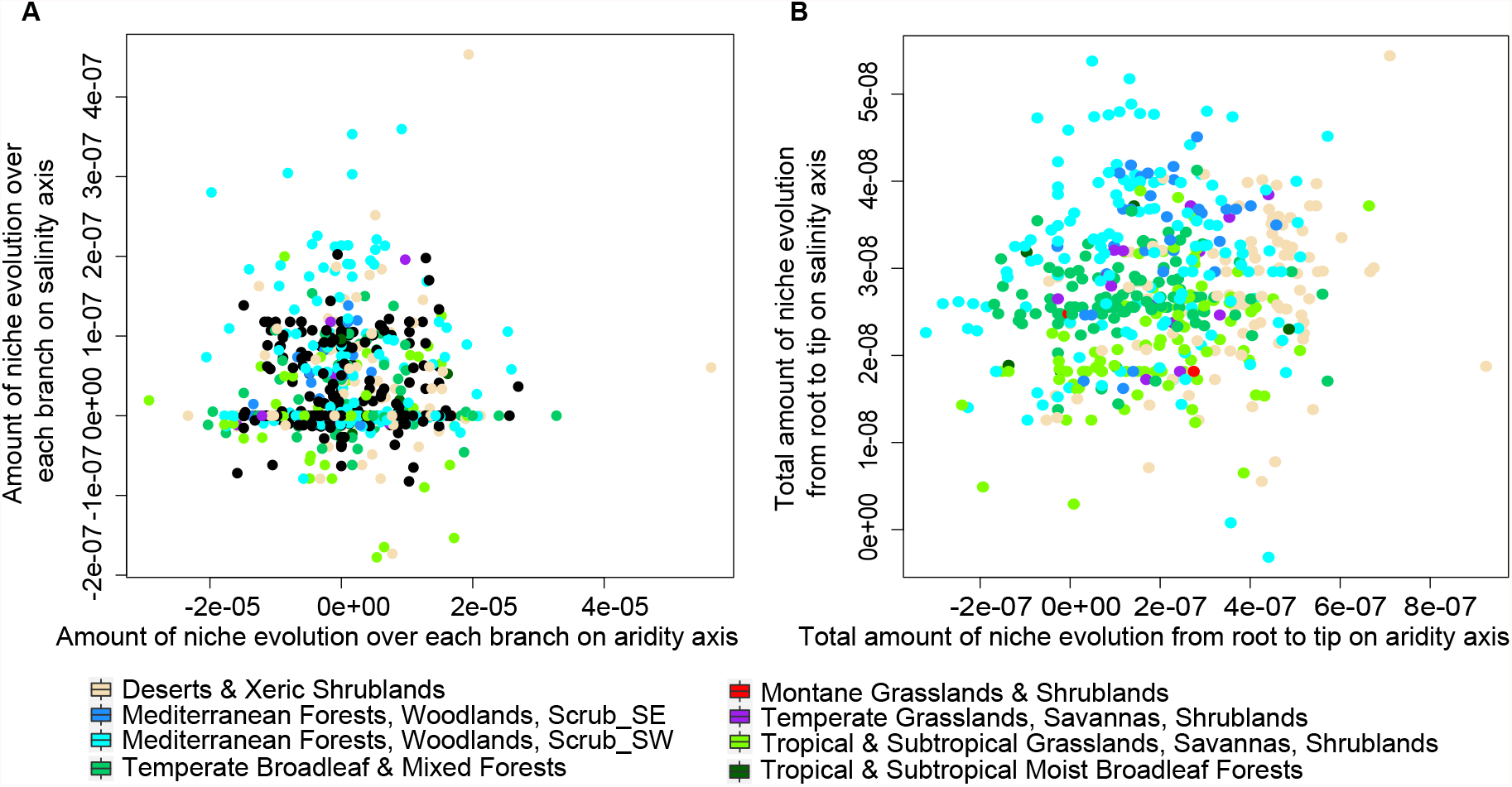
Correlation in the evolution of environmental tolerance along aridity and salinity. A) plots the amount of change in parameter value αof the fundamental niche long each branch. So each datapoint is a branch, with internal branches in black and terminal branches colored by the habitat that the majority of current presence location of the corresponding tip species fall in. B) plots the total amount of change in parameter value αof the fundamental niche from root to each tip species. So each datapoint is a tip species, colored by the habitat that the majority of the species’ presence location fall in. The plot for species in each habitat is in Figure S6. The same color scheme is used for habitats as Figure S1.

## Discussion

We reconstructed the evolution of the widespread plant genus *Acacia* over time and across different habitat types and infer a preponderance of evolutionary change towards more extreme values of aridity and salinity, inferring that the *Acacia* common ancestor could tolerate medium aridity level and very low salinity level, compared to contemporary levels in Australia. Our results suggest that there are multiple rounds of evolution of environmental tolerance in the ancestors of species currently found in arid and Mediterranean regions in both the old clades and the recent radiations. In contrast, we infer relatively little increase in aridity in recent radiations that lead to present-day species in tropical and subtropical forests and grasslands. Our results suggest different tempo and mode of evolution of aridity and salinity tolerance: while adaptation events increasing aridity tolerance are more frequent during the major periods of aridification in Australia, adaptation to higher salinity occurs over all periods. However, both salinity and aridity tolerance are inferred to have undergone a burst of adaptation events on both axes in the Pleistocene. These differing patterns suggest a decoupling of evolution of aridity and salinity tolerance, even though the tolerance level of aridity and salinity correlates in present-day species. As we explain below, these results are consistent with the adaptation of *Acacia* lineages to drier climates, allowing them to exploit new environmental opportunities opening up through the aridification of the continent. Our results are consistent with evolutionary flexibility in *Acacia* that may have played a key role in their current near-ubiquitous distribution across Australia and their dominance of many habitats.

The preponderance of evolution towards more extreme values of both aridity and salinity suggests that aridification has been a dominant driver of *Acacia* evolution and diversification across Australia during the Neogene and Quaternary. The multiple events of niche evolution to tolerate higher aridity and salinity on branches leading to species currently found in more arid and salt-affected regions suggest strong adaptation ability of *Acacia* to tolerate extreme environments. MRCAs of recent radiations in tropical and subtropical forests and grasslands can tolerate the same level of aridity as the common ancestor of all *Acacia* species included in the phylogeny. Since the common ancestor of *Acacia* already had the ability to tolerate the aridity level typical of non-desert areas in today’s Australia, our results support the hypothesis proposed by Renner et al. (2020) that *Acacia* was able to replace many other species groups in these mesic areas during the drying period of the Pleistocene, because these areas had increased aridity during the drying period and *Acacia* tolerated higher aridity. We also find a burst of adaptation events on both aridity and salinity axes in the Pleistocene, particularly in these recent radiations. This enhanced rate of niche evolution was also found by Renner et al. (2020) using traditional PCMs, which hypothesized that Pleistocene glacial-interglacial climate cycling (< 2.5Ma) drove rapid climatic adaptation. Yet, we find no evidence that niche evolution along aridity and salinity axes promoted diversification and led to the radiations, as Renner et al. (2020) hypothesized, because we find that the occurrence density of speciation events that are associated with niche evolution is not elevated during the Pleistocene. This result does not rule out the possibility that diversification is promoted by niche evolution along environmental axes other than aridity or salinity, but our analyses show that a clade with both high diversification rate and high rate of niche evolution does not necessarily suggest that niche evolution promotes diversification.

Comparing salt tolerance to aridity tolerance in *Acacia*, we find that the amount of evolution along aridity and salinity axes was not correlated across branches of the *Acacia* phylogeny. However, our inferred fundamental niches of present-day species are correlated between their tolerance abilities to aridity and salinity. This result is consistent with the observation that saline soils are frequently associated with aridity, so species need to adapt to both aridity and salinity to survive (Gong et al. 2017). Traditional PCMs based on the distribution data of tip species have been applied to test correlated evolution between different types of tolerance in *Acacia*. For example, correlated evolution was suggested between salinity and alkalinity tolerance in *Acacia* using phylogenetically independent contrasts (Bui et al. 2014b). Since traditional PCMs often assume Brownian motion evolution, they often infer correlated evolution if aspects of tip realised niches are correlated. Our analysis show that even when tip niches on different axes are correlated, we cannot assume that niche evolution along these axes occurs in concert, simply because there could be different histories to generate similar tip niches.

It is important to note that for the NEMo analysis, as for most macroevolutionary analyses, we have assumed that the phylogeny has accurate topology and provides an approximate scaling of evolutionary events in time. Clearly, phylogenetic estimates of either topology or the timing of evolutionary events are not error-free (Bromham et al. 2018; Guindon 2020; Nie et al. 2019). In particular, uncertainty or bias in branch length estimates arising from imperfect characterisation of molecular rates over time may have a non-trivial impact on macroevolutionary inference (Duchêne et al. 2017). Phylogenies such as this one where the only calibration is a root date may be particularly vulnerable to errors in position of node heights in some parts of the tree (Duchêne et al. 2014). One possible solution is to apply the NEMo analysis on posterior samples of phylogenies, but perhaps a better advance will be to embed the NEMo method into phylogenetic reconstruction, which allows us to directly link evolution on environmental tolerance to diversification events.

Another issue with our current inference of evolution in *Acacia* is incomplete sampling, as the *Acacia* phylogeny only includes half of the known extant species, and sampling is not random either with respect to area or lineage (Renner et al 2020). One obvious solution is to sample more species, but another way to increase the amount of information in the analysis is to incorporate historical data, such as paleoclimate or fossil distribution data, to supplement current species distribution data. For example, we could adjust priors for the occurrence rate of adaptation events with increasing aridity to allow higher rate under the known drying periods in Australia. Or we could include fossil locations and reconstructed paleoclimate by using the ENM equations to calculate the probability of sampling fossils at the locations given its inferred niches on the phylogeny and incorporate the probability into the posterior probability of NEMo. Here we have deliberately used only part of the available information in the inference process, so we can compare our inferred evolutionary history to known paleoclimatic history in order to demonstrate the utility of the method, but the method allows for all available information to contribute to stronger inference.

## Conclusion

To understand how plants in today’s Australia have responded to continuing aridification through geological time, we use a newly developed approach to reconstruct the evolution of tolerance to increasing aridity and soil salinity in a major Australian plant genus *Acacia*. By explicitly modelling changes in the fundamental niche, realised niche, and available niche of *Acacia* along its phylogeny, we find evidence that is consistent with *Acacia* having great adaptability to increasing aridity and salinity and that *Acacia*’s ability to tolerate aridity allows it to dominate not only the arid areas, but also the mesic areas by surviving the drying periods during Pleistocene in Australia. But unlike previous studies, we find no evidence that *Acacia*’s adaptability to increasing aridity and salinity promoted its radiation in today’s mesic areas. Neither do we find evidence for correlated evolution between tolerance to aridity and salinity, even when present-day *Acacia* species often need to adapt to both high aridity and high soil salinity. These results demonstrate the value of modelling biologically-informed processes of niche evolution, because we cannot make reliable inferences on processes from patterns without modelling the processes.

## Acknowledgements

We thank Joe Miller for providing *Acacia* distribution data, and we are grateful to Joe Miller, Elisabeth Bui, John La Salle, Rebecca Pirzl, and Lee Belbin for sharing their insight and expertise with us.

## Biosketch

Xia Hua is a mathematical biologist who specialises in finding analytical solutions to a wide range of evolutionary problems, including research in molecular evolution, macroevolution and macroecology and language evolution. Marcel Cardillo’s research aims to understand large-scale patterns of biodiversity, with a current focus on Australian plants. Lindell Bromham designs comparative analyses for investigating mechanisms for the generation of diversity, from molecular evolution to macroevolution to languages and cultures. Together, they are the three principal investigators from the Macroevolution and Macroecology Group at the Australian National University.

## Editor

Margaret Byrne

## Data availability

This study uses publicly available data on *Acacia* phylogeny and distribution, from sources cited in the text. Code with step-by-step instructions for implementing the model can be found at https://github.com/huaxia1985/NEMo.

