## Supplementary Materials for "Adapting to extremes: reconstructing evolution in response to changing climate over time and space in the diverse Australian plant genus *Acacia*"

**Supplementary Methods:**

1. Accounting for sampling bias
2. Model Parameters
3. MCMC approximation

**Figure S1.** Phylogenetic distribution of the inferred history of niche evolution with highest posterior probability along axes of aridity and salinity.

**Figure S2.** Trends in posterior probabilities over generations for the 20 rjMCMCs in aridity analysis and salinity analysis.

**Figure S3.** The amount of evolution in fundamental niche per branch in each posterior sample of the 20 rjMCMCs after burnin in aridity analysis.

**Figure S4.** The amount of evolution in fundamental niche per branch in each posterior sample of the 20 rjMCMCs after burnin in salinity analysis.

**Figure S5.** Occurrence density of the inferred speciation and adaptation events per branch in each major clade of the phylogeny that fall within each time interval.

**Figure S6.** Correlation in the inferred fundamental niche of tip species with majority presence locations fall in a habitat type.

### *1. Accounting for sampling bias*

We used an inhomogeneous Poisson point process (Renner et al. 2015) as the ENM component of NEMo. Because opportunistic sampling bias can influence the niche estimation (Fithian et al. 2015), we modified the likelihood function to account for sampling bias, using the method of Fithian et al. (2015). The log-likelihood of the presence-only data of each tip species $I_{PO}$ is modified as,

$$\ell_P = \sum_{i \in I_{PO}} \left[\log\left(\lambda_{r_i}\right) + \delta z_i\right] - \frac{\mathcal{D}}{N} \sum_{i \in I_{BG}} \lambda_{f_i} \lambda_{a_i} e^{\delta z_i}$$

, and the log-likelihood of the presence-absence data of each tip species $I_{PA}$ is

$$\ell_A = \sum_{i \in I_{PA}} \left[-\mathrm{y}_i \log\left(1 - e^{-\lambda_{r_i}}\right) + (1 - y_i)\lambda_{r_i}\right]$$

, where $\mathrm{y}_i = 1$ if the species is present at the $i^{\mathrm{th}}$ location and $\mathrm{y}_i = 0$ if the species is absent at the $i^{\mathrm{th}}$ location. For both equations, $\lambda_{r_i}$ is the realised niche, or the probability of recording a presence of the species under the environmental condition at the $i^{\mathrm{th}}$ location. For presence-only data, $z_i$ are values of the factors that influence sampling effort at the $i^{\mathrm{th}}$ location and $\delta$ are the coefficients of the two sampling factors we included - distance of the location to the nearest road and whether the location is in a protected area - which are assumed to have the same value for all tip species. The starting values of coefficients were set to their maximum likelihood estimates given the likelihood function, and a rather flat prior was used: an exponential distribution with rate of 0.1. $\lambda_f$ is the inferred fundamental niche of a tip species and $\lambda_a$ is the inferred available niche of the tip species. If we were not accounting for sampling bias, the second term of the equation would be a direct integral of $\lambda_f \lambda_a$ (Hua et al. 2021), but because each location now has a specific value for sampling bias, we can only approximate the integral by randomly sampling $N$ locations from the surroundings of the species presence locations to construct a set of background locations $I_{BG}$

and averaging the term to integrate over all the background locations. Note that these background locations are only used to numerically approximate the second term of the equation, so they are not pseudo-absence locations (Fithian et al. 2015).

For each *Acacia* species, background locations were sampled uniformly from areas within an arbitrary large radius (20 $km$) of each presence location of the species, so the weight of each background location is $\frac{\mathcal{D}}{N}$, where $\mathcal{D}$ is the total area that falls within the 20 $km$ radius of each species presence location, and $N$ is the total number of background locations, which was ten times the number of the species presence locations, with a minimum of 1000 for species with few presence locations. Based on the environmental conditions over all the background locations, we used Sturges' formula (Sturges 1926) to decide the bin size to discretise the aridity index and the electrical conductivity.

### *2. Model Parameters*

NEMo has four universal parameters which are assumed to be constant across the phylogeny: heritability, dispersal rate, the occurrence rate of adaptation events, and the occurrence rate of speciation events. The starting values for heritability and dispersal rate were arbitrarily set to 0.03 and 1, respectively. Dispersal rate had a rather flat prior, an exponential distribution with rate of 0.5. Heritability had a narrow prior, an exponential distribution with rate of 20. The narrow prior for heritability was used because parameters of the fundamental niche change proportionally to heritability on a log scale. The starting value for the occurrence rate of adaptation event and the occurrence rate of speciation event was arbitrarily set to 0.1. A narrow prior was applied on both occurrence rates, an exponential distribution with rate of 10 to reduce the degree of overestimation on the number of events (see Hua et al. 2021). Additional parameters that are specific to each adaptation and speciation event are: 1) amount of environmental change in an adaptation event, 2) proportion of a tolerance range that is inherited by one descendent species (its sister species inherits the other part) from the ancestral species in a speciation event, 3) where a dispersal barrier locates along the environmental axis. Uniform priors were used for these parameters (see detailed explanation in Hua et al. 2021).

We apply separate NEMo analyses to the evolution of environmental tolerances in *Acacia* along aridity and salinity axes. For each of these axes, the starting values for parameters of root niches and their prior means were set to their maximum likelihood estimates given the probability of each tip species' presence and/or absence data, assuming that no events of niche evolution occurred on the phylogeny. The prior for mean of the root available niche had a normal prior with standard deviation of 3. The prior for the standard deviation of the root available niche had a lognormal prior with logarithmic standard deviation of 0.5. The prior for the parameters on log-scale for the root fundamental niche ($\alpha$ and $\beta$) had normal priors with standard deviation of 1.

***3. MCMC approximation***

For both aridity and salinity, we ran 20 independent reversible-jump Markov Chain Monte Carlo (rjMCMCs) for $3\times10^6$ generations, with a thinning interval of 1000. The 20 rjMCMCs of aridity analysis converge to similar posterior probabilities, while 20 rjMCMCs of salinity analysis do not converge (Figure S2). This may suggest that current distributions of tip species are not informative to the evolutionary history of salt tolerance in *Acacia*, compared to the tolerance to aridity. This is also suggested in the posterior distribution of the heritability that the heritability of salinity tolerance is 1/100 to 1/1000 of the heritability of aridity tolerance. Note that heritability is on an evolutionary time scale, so low heritability equates high phylogenetic lability (high rates of gain and loss) in salt tolerance, which has been suggested by different analyses in grasses (Bennett et al. 2013; Moray et al. 2015; Bromham et al. 2015).

However, for both analyses, the 20 rjMCMCs agree on a general pattern of the amount of change in the fundamental niche along *Acacia* phylogeny (Figure S3 and S4). The reason why rjMCMCs of salinity analysis do not converge but agree on a general pattern of niche evolution is that each tip species has many presence/absence locations and tip species have more narrow and more different realized niche to each other along salinity than aridity, so small change in the local parameters of a single adaptation or speciation event can cause large change in the posterior probability.

After discarding the first half of the generations as burnin, global parameters other than root niche parameters have large effective sample size over 450. Root niche parameters, which include the mean and standard deviation of root available niche and $\alpha$ and $\beta$ of root fundamental niche, have relatively low effective sample size between 50 to 200. This is due to their correlation with local parameters of adaptation and speciation events, which reduces sampling efficiency of rjMCMC. For example, a root with low tolerance level plus adaptation events at all the basal branches would give similar posterior probability to a root with high tolerance level without adaptation events at the basal branches. So a big change in root tolerance level is only sampled when there is a big change in the adaptation events at the basal branches. However, this should not affect our inference of the tolerance level at basal branches, which is robust to the root niche parameters. Whether the root niche parameters are for a medium or low aridity level, there are adaptation events inferred along the basal branches that increase the tolerance level of the basal branches, suggesting that the early ancestors *Acacia* lineage had tolerance level to medium aridity level compared to today's Australia.

**Figure S1.** Phylogenetic distribution of the inferred history of niche evolution with highest posterior probability along axes of aridity (left) and salinity (right). *Acacia* is a diverse genus of trees and shrubs occupying many habitats across Australia: for example, while silver wattle *A. dealbata* (A) is found in relatively mesic habitats in southern Australia, waddi trees *A. peuce* (B) grow in arid conditions in central Australia. *Acacia* are found across diverse ecoregions in Australia, shown in panel (C) using the World Wildlife Fund's categories of terrestrial ecoregions. Spatial variation in the aridity index in today's Australia is shown in (D), with higher values indicating more arid areas. (E) shows the variation in root zone soil electrical conductivity in today's Australia, with higher values indicating areas with higher salinity. In the phylogeny, each tip species has a stacked bar showing the relative proportion of its current presence locations in each habitat, in the same color scheme as the habitat map. The blue to red branch color gives the amount of change in the parameter value $\alpha$ of the fundamental niche long the branch. The pattern in parameter value $\beta$ is similar, so, for illustration, we only plot α. Red indicates tolerance to more extreme conditions and blue indicates tolerance to less extreme conditions. The inferred realised niche for each tip and each node that has large change in realised niche from its ancestral node is shown in a row of grids. Each grid corresponds to an environmental condition that becomes more extreme from left grid to right grid, and grid color gives the relative occurrence probability of the tip or node under each environmental condition. The inferred speciation (circle) and adaptation (triangle) events are plotted at the location where it occurred on the phylogeny, with the blue to red color giving their corresponding local parameter value. For a speciation event, red indicates the branch inherits the part of realised niche under more extreme conditions, and blue indicates the branch inherits the part under less extreme conditions. For an adaptation event, red indicates environmental change to more extreme conditions and blue to less extreme conditions. We also label major clades in the phylogeny, which corresponds to the clade label in Figure S3-S5. Image of *A. dealbata* by SABENCIA Bertu Ordiales (reproduced under Creative Commons Attribution-Share Alike 4.0 International license). Image of *A. peuce* by Marcel Cardillo.

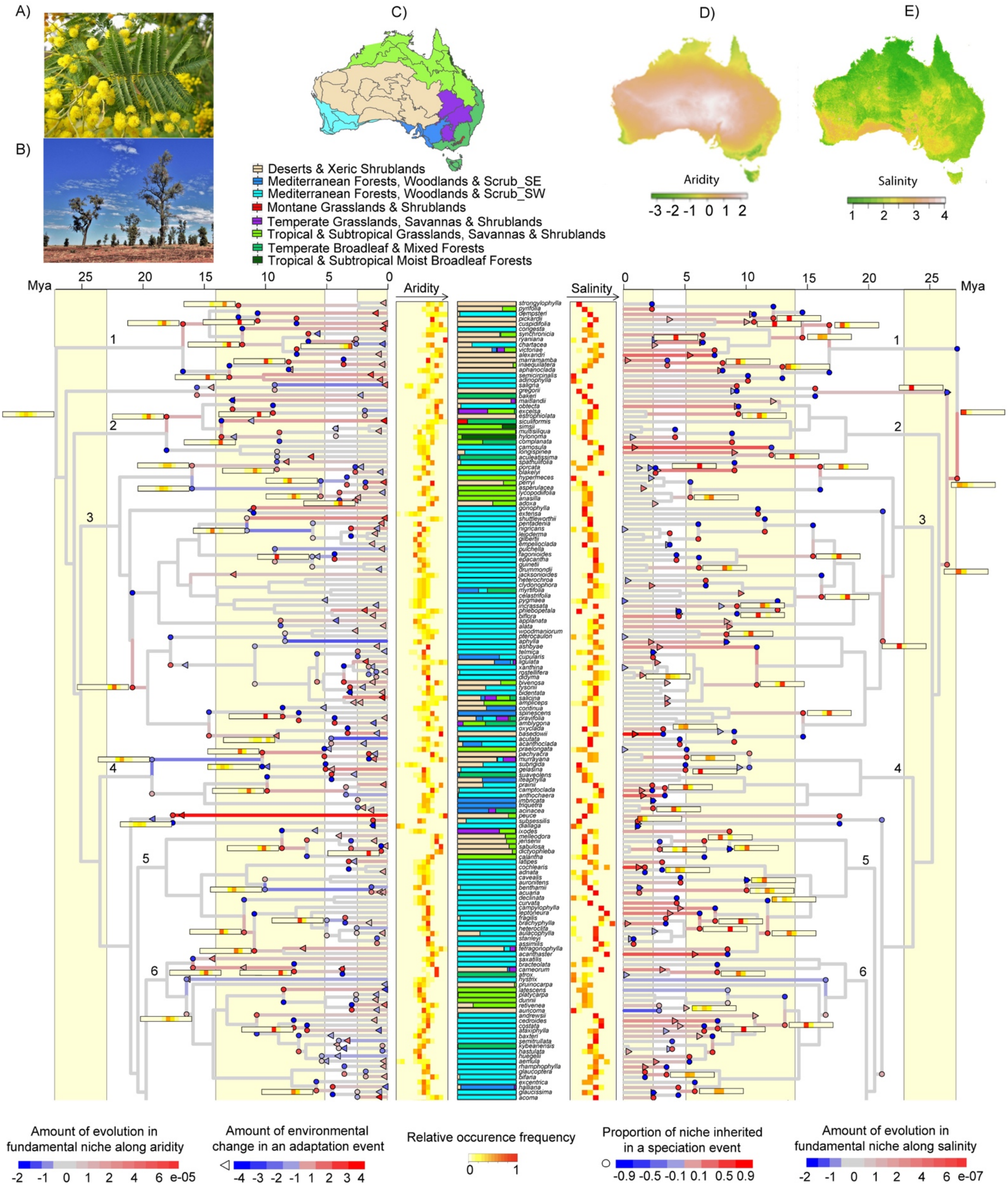

A)
B)
C)
D)
E)
Deserts & Xeric Shrublands
Mediterranean Forests, Woodlands & Scrub_SE
Mediterranean Forests, Woodlands & Scrub_SW
Montane Grasslands & Shrublands
Temperate Grasslands, Savannas & Shrublands
Tropical & Subtropical Grasslands, Savannas & Shrublands
Temperate Broadleaf & Mixed Forests
Tropical & Subtropical Moist Broadleaf Forests
Aridity
-3 -2 -1 0 1 2
Salinity
1 2 3 4
Mya
25 20 15 10 5 0
Aridity
Salinity
0 5 10 15 20 25
Mya
1
2
3
4
5
6
Amount of evolution in fundamental niche along aridity
-2 -1 0 1 2 4 6 e-05
Amount of environmental change in an adaptation event
-4 -3 -2 -1 1 2 3 4
Relative occurence frequency
0 1
Proportion of niche inherited in a speciation event
-0.9 -0.5 -0.1 0.1 0.5 0.9
Amount of evolution in fundamental niche along salinity
-2 -1 0 1 2 4 6 e-07

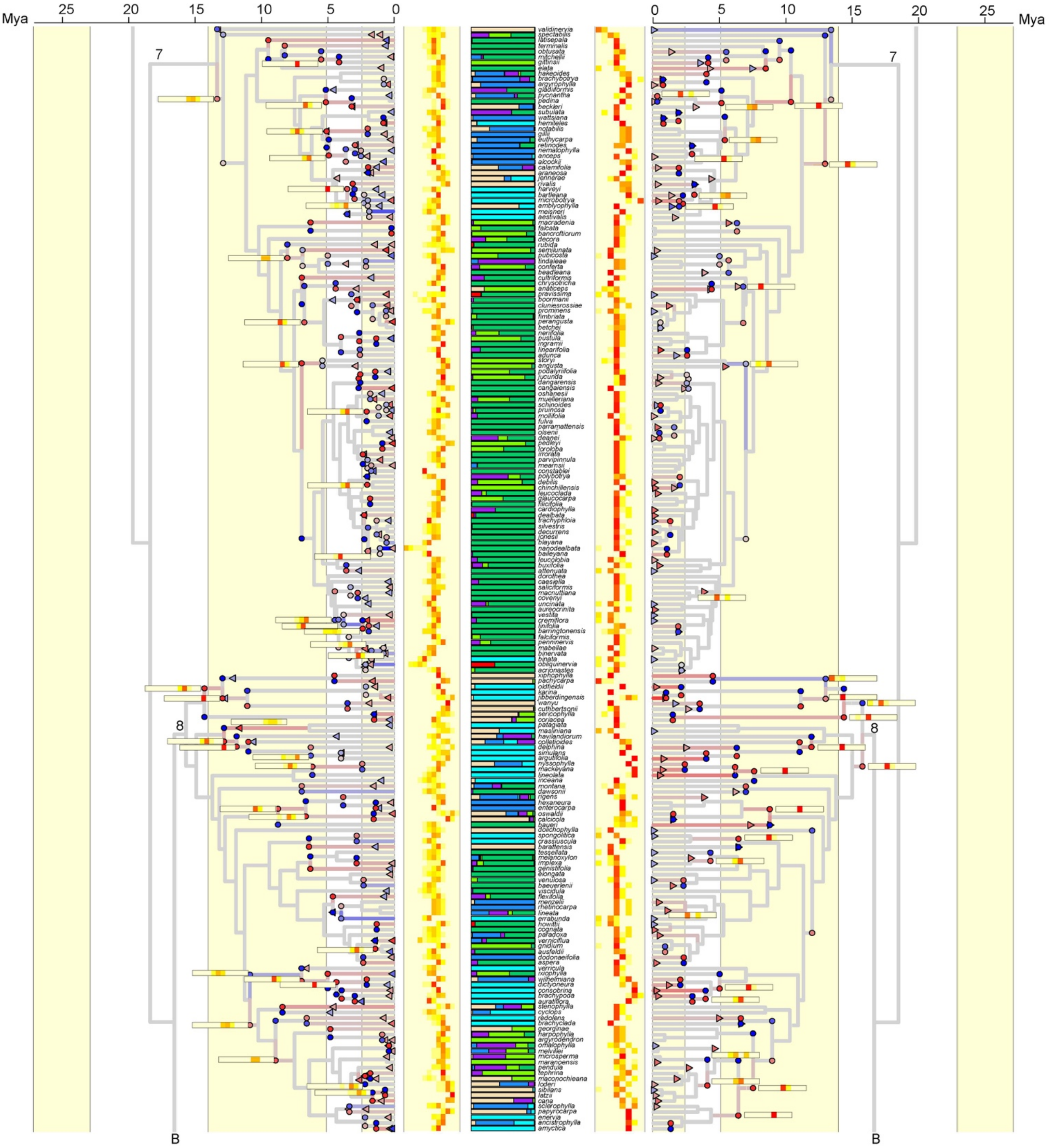

Mya
25
20
15
10
5
0
7
8
B

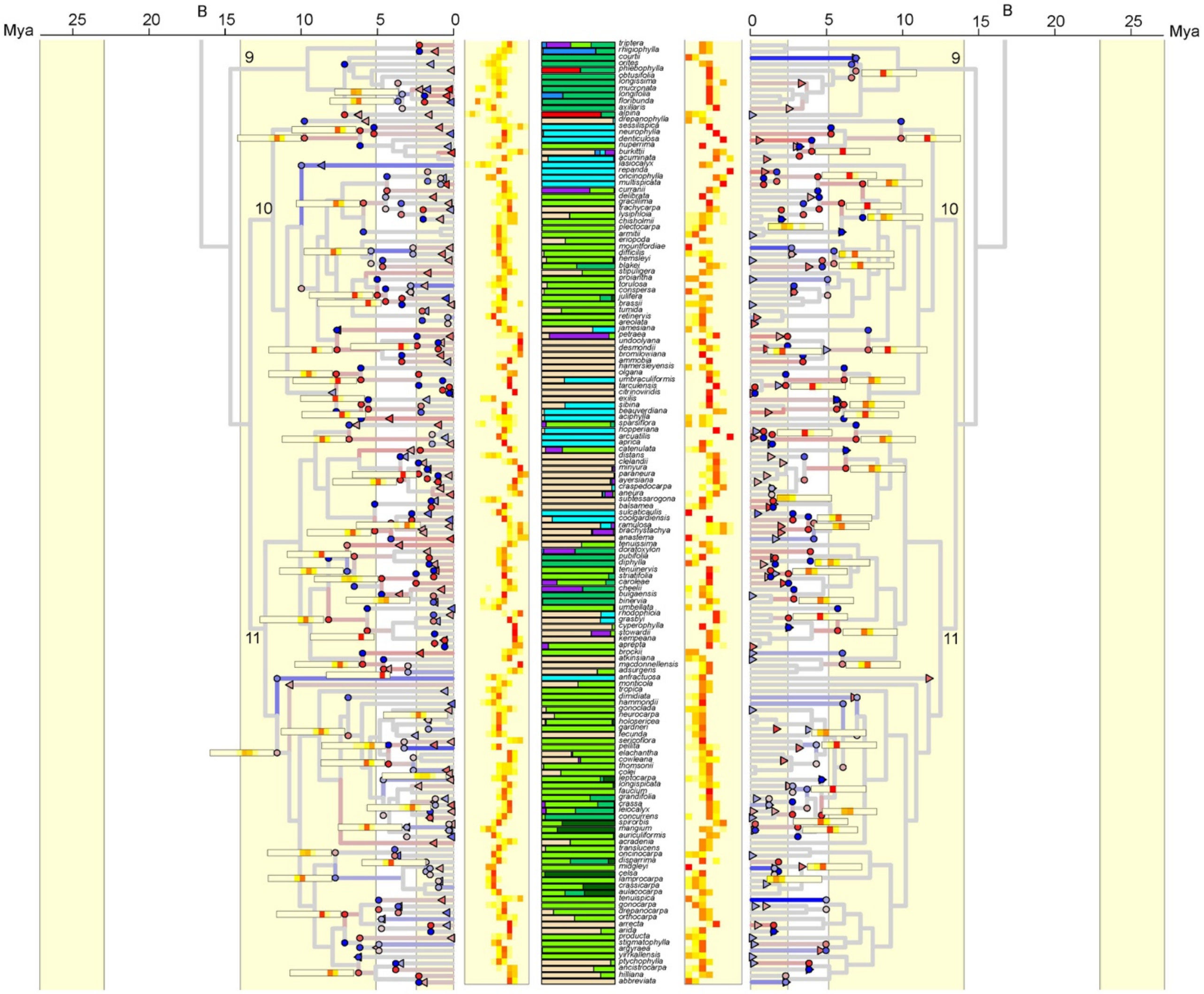

Mya
25
20
B
15
10
5
0
9
10
11

**Figure S2.** Trends in posterior probabilities over generations for the 20 rjMCMCs in aridity analysis and salinity analysis. Each line is an rjMCMC run.

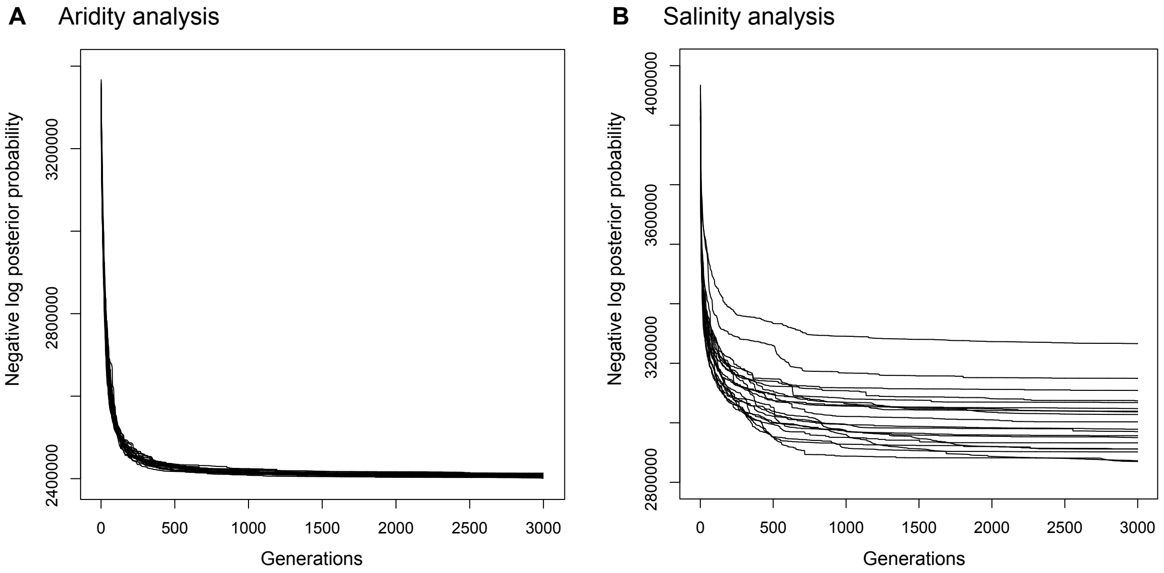

**Figure S3.** The amount of evolution in fundamental niche per branch in each posterior sample of the 20 rjMCMCs after burnin in aridity analysis. Each column is an rjMCMC. Within each column, each horizontal bar corresponds to a branch in the phylogeny and is consist of 1500 grids, each corresponding to a posterior sample after burnin. Each grid is colored by the amount of change in the parameter value $\alpha$ of fundamental niche along a branch. The fact that these grids have such a similar color that they connect to a bar suggests that posterior samples of the same rjMCMC converge to the same history of niche evolution on the phylogeny. To help identify the corresponding branch in the phylogeny, internal branches are grouped separately from external branches, and branches or bars that are from the same clade is grouped together with the clade label same as in Figure 1. Comparing columns, we can see a general pattern in the inferred history of niche evolution among rjMCMCs. For example, none of the rjMCMCs infer niche evolution on the basal branches in the phylogeny.

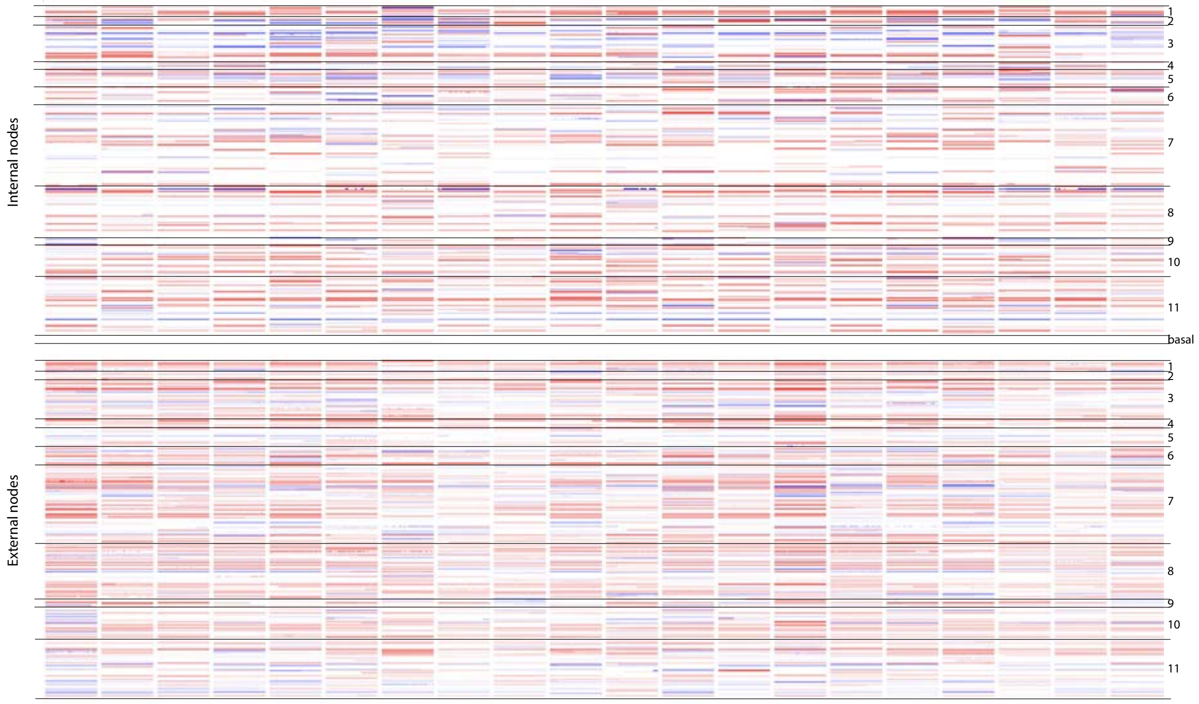

**Figure S4.** The amount of evolution in fundamental niche per branch in each posterior sample of the 20 rjMCMCs after burnin in salinity analysis. Each column is an rjMCMC. Within each column, each horizontal bar corresponds to a branch in the phylogeny and is consist of 1500 grids, each corresponding to a posterior sample after burnin. Each grid is colored by the amount of change in the parameter value $\alpha$ of fundamental niche along a branch. The fact that these grids have such a similar color that they connect to a bar suggests that posterior samples of the same rjMCMC converge to the same history of niche evolution on the phylogeny. To help identify the corresponding branch in the phylogeny, internal branches are grouped separately from external branches, and branches or bars that are from the same clade is grouped together with the clade label same as in Figure 1. Comparing columns, we can see a general pattern in the inferred history of niche evolution among rjMCMCs. For example, all rjMCMCs infer increasing tolerance to salinity on the basal branches in the phylogeny.

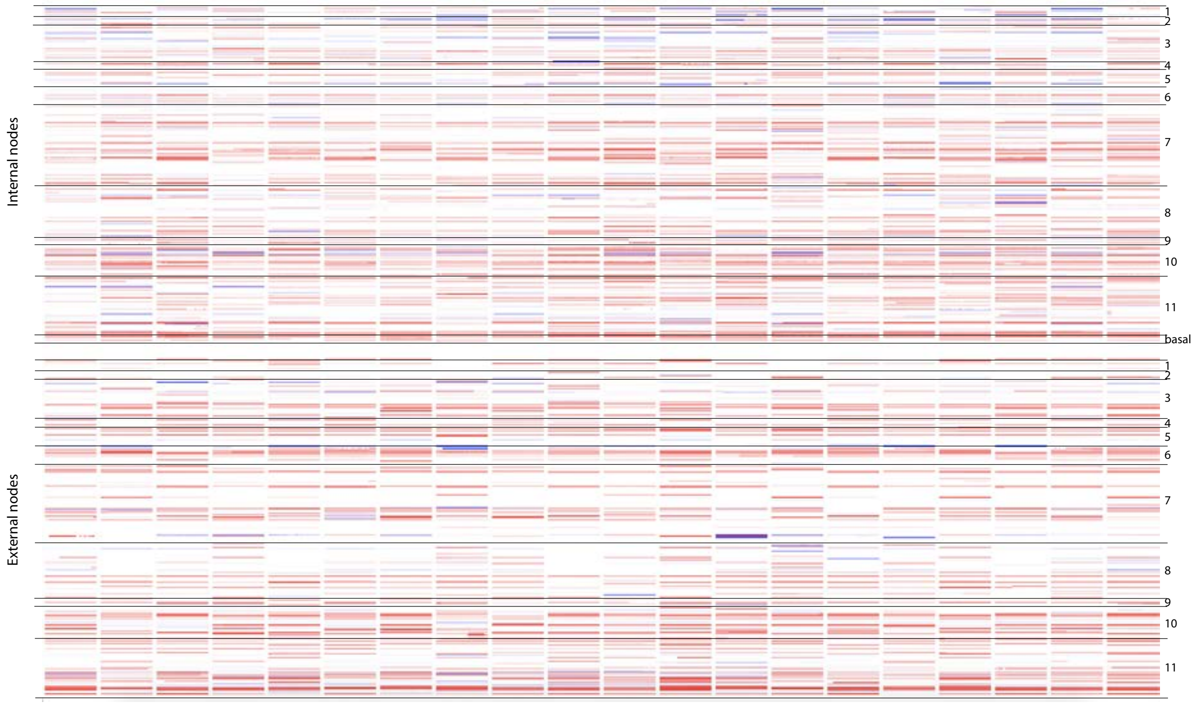

**Figure S5.** Occurrence density of the inferred speciation and adaptation events per branch in each major clade of the phylogeny that fall within each time interval. This figure is similar to Figure 4, but with each plot for tip species in the same clade of the phylogeny. Clade label is the same as in Figure 1.

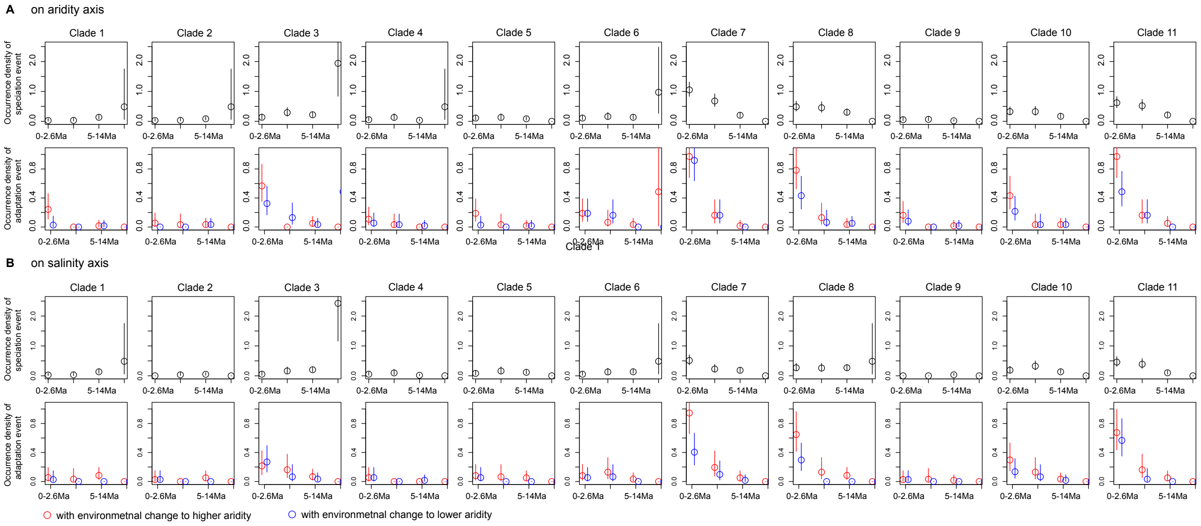

**Figure S6.** Correlation in the parameter value $\alpha$ of the fundamental niche inferred for each tip species that have majority presence locations fall in a habitat type. Montane habitat is not plotted because there are only two montane species.

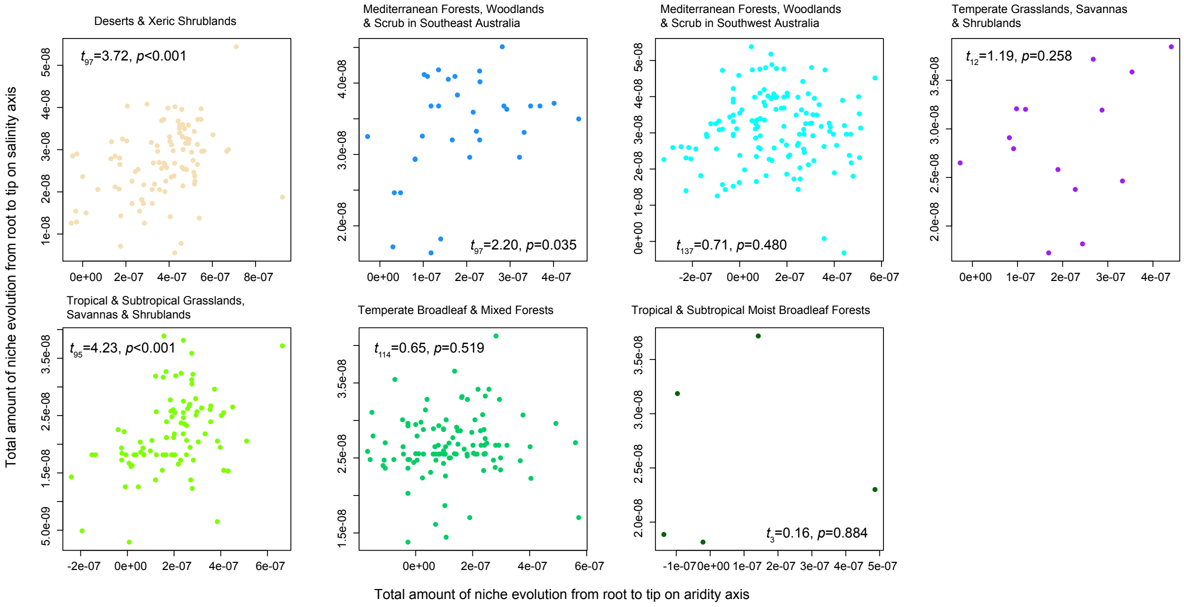
